# Identifying Neural Correlates of Balance Deficits in Traumatic Brain Injury Using Partial Least Squares Correlation Analysis

**DOI:** 10.1101/2022.05.15.491997

**Authors:** Vikram Shenoy Handiru, Easter S. Suviseshamuthu, Soha Saleh, Haiyan Su, Guang H. Yue, Didier Allexandre

**Author notes:** Correspondence should be addressed to Vikram Shenoy Handiru.

## Abstract

**Background:** Balance impairment is one of the most debilitating consequences of Traumatic Brain Injury (TBI). To study the neurophysiological underpinnings of balance impairment, the brain functional connectivity during perturbation tasks can provide new insights. To better characterize the association between the task-relevant functional connectivity and the degree of balance deficits in TBI, the analysis needs to be performed on the data stratified based on the balance impairment. However, such stratification is not straightforward, and it warrants a data-driven approach.

**Approach:** We conducted a study to assess the balance control using a computerized posturography platform in 17 individuals with TBI and 15 age-matched healthy controls. We stratified the TBI participants into balance-impaired and non-impaired TBI using *k* -means clustering of either center of pressure (COP) displacement during a balance perturbation task or Berg Balance Scale (BBS) score as a functional outcome measure. We analyzed brain functional connectivity using the imaginary part of coherence across different cortical regions in various frequency bands. These connectivity features are then studied using the mean-centered partial least squares correlation (MC-PLSC) analysis, which is a multivariate statistical framework with the advantage of handling more features than the number of samples, thus making it suitable for a small-sample study.

**Main Results:** Based on the nonparametric significance testing using permutation and bootstrap procedure, we noticed that the theta-band connectivity strength in the following regions of interest significantly contributed to distinguishing balance impaired from non-impaired population, regardless of the type of strat-ification: *left middle frontal gyrus, right paracentral lobule, precuneus*, and *bilateral middle occipital gyri*.

**Significance:** Identifying neural regions linked to balance impairment enhances our understanding of TBI-related balance dysfunction and could inform new treatment strategies. Future work will explore the impact of balance platform training on sensorimotor and visuomotor connectivity.

## 1. Introduction

Traumatic Brain Injury (TBI) is one of the leading causes of death and disability worldwide [1]. With the immediate consequences of long-term disability due to injury, individuals with TBI are often at an elevated risk of impaired motor functions, such as loss of postural control [2]. According to a longitudinal follow-up evaluation, approximately 60% of persons with TBI exhibit persistent balance deficits two years after the injury [3]. Depending on the anatomical extent of TBI, it can significantly damage one or more of the key components of the postural control system. These damages due to TBI can manifest in different ways, such as vestibular dysfunction, postural hypotension, sensory deficits, and brainstem injury [4].

Previous structural brain mapping studies have shown that neural insults to the white matter tracts in the thalamus, corpus callosum, and corticospinal tracts can disrupt cross-hemispheric communication in the brain [5]; [6]. Relevant research in functional brain mapping has also shown that altered global functional network organization can lead to balance deficits due to the disruption of sensorimotor, execution, and control networks [7]. However, the task-relevant neurophysiological mechanisms leading to the postural imbalance are still unknown [8].

It is worth noting that one of the limitations of balance-related functional brain mapping, as highlighted by [9], is that the data is often collected independently from a performance-based balance function assessment of individuals with TBI. For example, most fMRI studies fall into this category since it is not possible to assess the neural correlates of standing postural deficits. Past fMRI-based studies have investigated resting-state fMRI and its association with physical balance measures [10]; [11]. In this regard, EEG is an attractive option for studying postural task-relevant functional brain mapping even in individuals with TBI [12]; [13]. The limitation, however, is that these studies do not explain task-relevant functional connectivity. With the premise that the postural imbalance can be attributed to the loss of sensorimotor integration [14], we propose that the neural correlates of balance deficits are better characterized using the functional connectivity between the regions involved in postural control tasks than using the cortical activity features. The specific mechanisms of balance control in terms of the sensory organization are presented in [15]. Although there are some studies evaluating balance-related brain functional connectivity changes in healthy individuals using fNIRS [16] and EEG [17], there are very few studies on individuals with TBI [18]; [19].

To address research questions concerning balance deficits in TBI, it is essential to objectively stratify the TBI population into groups with and without balance impairments, recognizing that balance deficits vary among TBI patients. Studies have attempted this stratification [10]; [20]; [21], but with certain limitations. In [22], the stratification of individuals with TBI into those with and without balance deficits was based on the subjective (self-reported) questionnaire. Other studies investigating TBI individuals with and without balance deficits [20]; [21] did not study the neural mechanisms of the balance function. These challenges underscore the need to bridge the knowledge gap by examining brain dynamics in both balance-impaired and balance-non-impaired TBI populations in comparison to healthy controls.

Although a simple correlation analysis might initially seem appealing for investigating the link between brain activity or connectivity and quantifiable functional measures of balance deficits, the complexity of high-dimensional brain recordings and multiple functional outcomes necessitates corrections for multiple correlations, reducing the statistical power of the findings. Therefore, we choose partial least squares correlation (PLSC) that is well-suited for neuroimaging studies, and a mathematically well-founded and robust statistical approach for reliably predicting the relationship between a large set of variables despite a small sample size [23]. Moreover, it obviates the need for performing multiple comparisons—such as in the case of post-hoc analysis in the analysis of variance (ANOVA)—which is not a viable option with small samples. Interestingly, the PLS methods have recently been employed to identify the brain-behavior association, whereby higher connectivity between the default mode and sensorimotor network is associated with sport-related concussion and balance symptom severity [24]. Motivated by this multivariate data-driven approach, we employ the mean-centered PLS method (**Fig. 1**) to extract neural markers of balance impairment in chronic TBI. The input features for the PLSC are primarily the functional connectivity strength of different brain regions involved in the balance control. Moreover, our choice of features also aligns with the observation that the complex task of postural control, especially when subject to external perturbations, necessitates the engagement of multiple brain functional networks specific to each sub-task [25]. Together, our study data analysis goal is to introduce PLSC approach to determine which brain regions are crucial in distinguishing between balance-impaired and balance-non-impaired TBI patients.

**Figure 1:**
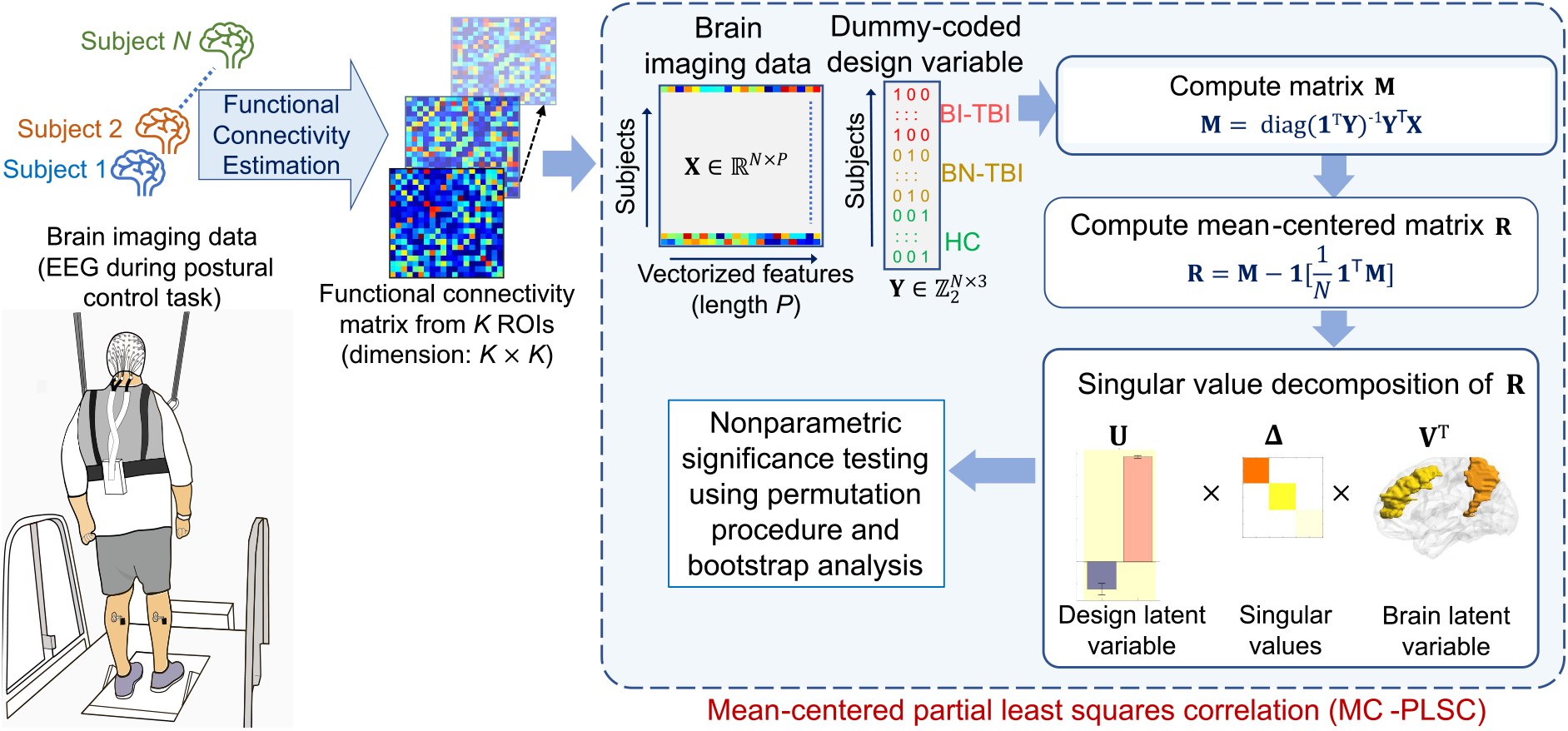
We performed the data analysis using the mean-centered partial least squares correlation (MC-PLSC) approach. The functional connectivity matrix involving K ROIs is constructed from the EEG data of each study participant. The design matrix **X** is built by row-wise stacking of vectorized functional connectivity matrices from N subjects. The dummy coding matrix **Y** is binary valued with N rows, and it represents the experimental groups. The group-wise average **M** is computed, and each column of **M** is subtracted from its mean value to obtain **R**. The singular value decomposition (SVD) of **R** yields the left and right singular vectors, denoted as **U** and **V**, respectively, and the corresponding singular values Δ . The singular vectors **U** and **V** are also known as design or contrast latent variable (LV) and brain LV, respectively. The statistical significance of the singular values and LVs are tested with the permutation procedure and bootstrap analysis, respectively.

## 2 Methodology

### 2.1 Participant Characteristics

This study enrolled 18 individuals with chronic traumatic brain injury (TBI) and 18 age-matched healthy controls (HC). Due to the very noisy EEG data of some participants, we had to exclude 1 TBI and 3 HC subjects from the analysis. Thus, this paper presents the data from 17 TBI and 15 HC. More information on the inclusion and exclusion criteria for the study participants can be found in our previously published research (Shenoy Handiru *et al* 2021). All the subjects provided a signed written informed consent before the experiment. The study was in accordance with the Declaration of Helsinki and approved by the Kessler Foundation Institutional Review Board (R-890-15).

The patient demographics and clinical characteristics of TBI participants are presented in Table 1.

**Table 1:** TBI Patient Demographics and Clinical Characteristics (N = 17)

| Subject ID | Gender | Age<br>(years) | Height<br>(cm) | Weight<br>(kg) | COP<br>(cm) | BBS | Severity | Years since injury |
| --- | --- | --- | --- | --- | --- | --- | --- | --- |
| ABI001 | M | 51 | 190.5 | 95.3 | 9.39 | 48 | Severe | 28 |
| ABI002 | M | 58 | 182.9 | 88 | 7.31 | 42 | Severe | 2 |
| ABI003 | M | 60 | 175.9 | 88.7 | 19.18 | 44 | Severe | 7 |
| ABI004 | M | 59 | 178.4 | 143.8 | 13.49 | 49 | Severe | 6 |
| ABI005 | M | 51 | 180.3 | 113.9 | 9.47 | 54 | Severe | 5 |
| ABI006 | M | 52 | 177.8 | 97.5 | 12.86 | 54 | Severe | 6 |
| ABI007 | F | 53 | 170.2 | 76.2 | 18.0 | 51 | Mild | 1 |
| ABI008 | M | 60 | 180.3 | 111.1 | 8.19 | 48 | Moderate | 1 |
| ABI009 | F | 32 | 167.6 | 58.5 | 10.12 | 56 | Mild | 3 |
| ABI010 | M | 49 | 172.7 | 66.7 | 14.14 | 46 | Moderate | 1 |
| ABI012 | F | 56 | 168.9 | 120.2 | 4.8 | 55 | Mild | 10 |
| ABI013 | F | 60 | 154.3 | 72.6 | 17.27 | 34 | Severe | 57 |
| ABI016 | M | 56 | 182.9 | 86.2 | 7.32 | 56 | Moderate | 6 |
| ABI018 | M | 35 | 182.9 | 68 | 15.33 | 44 | Moderate/severe | 9 |
| ABI019 | M | 23 | 177.8 | 63.5 | 7.44 | 55 | Severe | 3 |
| ABI020 | M | 50 | 187.96 | 102.1 | 9.17 | 55 | Moderate/severe | 10 |
| ABI021 | M | 23 | 182.88 | 89.4 | 14.54 | 52 | Moderate/severe | 2 |

**Table 2:**
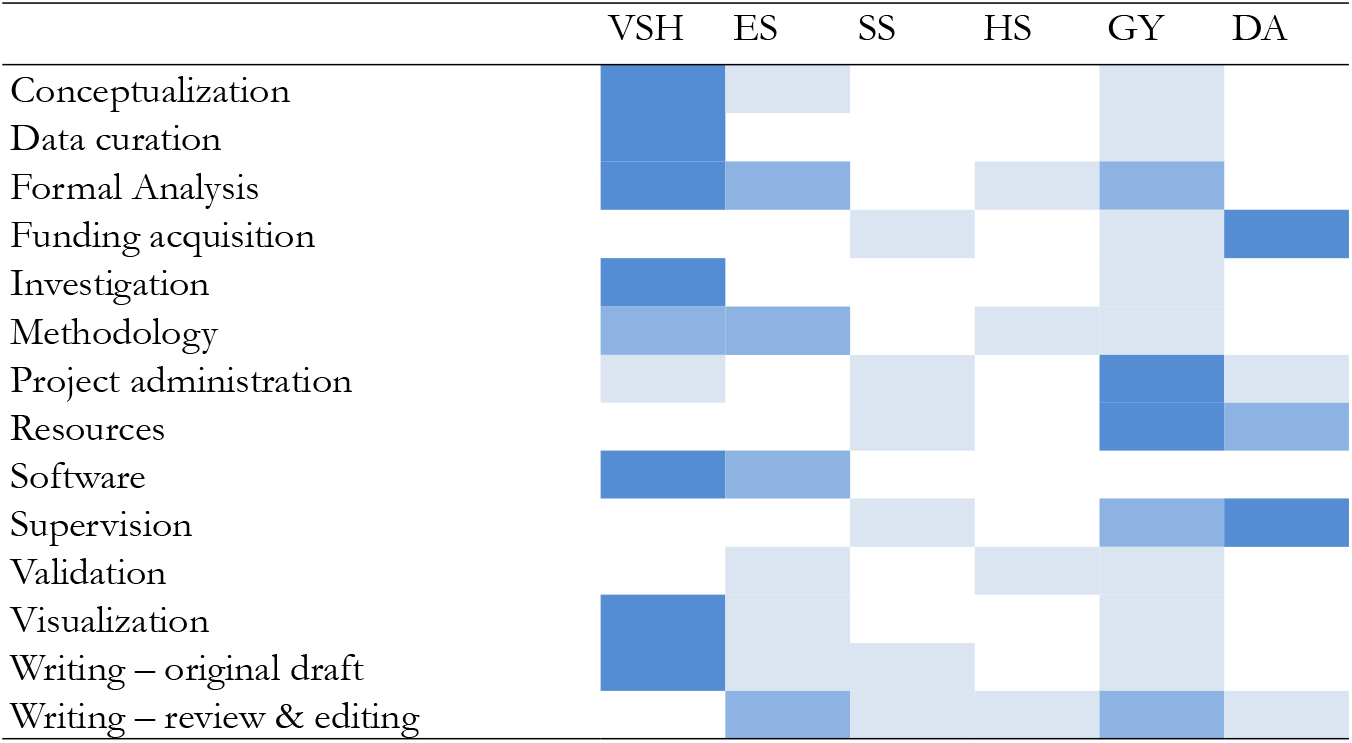
Authorship contribution listed according to the CRediT taxonomy.

### 2.2 Data Acquisition and Pre-processing

We used multiple modalities to collect EEG and the posturography platform data during a balance perturbation task and MRI to obtain subject-specific anatomical data for EEG source localization.

#### (A) MRI Data Acquisition

The MRI data (T1-weighted MPRAGE scan) was acquired at the Rocco Ortenzio Neuroimaging Center (Kessler Foundation, NJ) using the Siemens Skyra 3T scanner (Erlangen, Germany) with the following specifications: 1-mm isotropic voxel resolution, TE=3 ms, TR=2300 ms, 1-mm thick 176 slices, Field of View (FOV) 256x256 mm^2^.

#### (B) Posturography Data Acquisition

The perturbation-related data was recorded using the computerized dynamic posturography platform (NeuroCom Balance Master, NeuroCom Intl, Clackamas OR). This computerized platform was pre-programmed to generate unpredictable sinusoidal perturbations at low amplitude (0.5 cm) or high amplitude (2 cm) in the anterior-posterior (or forward and backward) direction at 0.5 Hz for 4 s with a random intertrial interval of 4-8 s. The posturography data were collected as 5 blocks, where each block consisted of 20 trials with high/low amplitude and forward/backward perturbation in a random order. We mainly presented the findings from a set of 29 trials of high amplitude backward perturbation, as it is the most challenging condition with the highest range of body sway across subjects.

#### (C) EEG Data Acquisition

The brain activities during the balance perturbation were noninvasively recorded using the 64-channel EEG system (ActiCAP BrainAmp standard, Brain Products®, Munich, Germany) at a sampling rate of 500 Hz. The EEG electrodes were positioned according to the extended 10-10 montage, with the electrodes FCz serving as the common reference and AFz as the ground. The electrode positions were digitized in 3D using the Brainsight Neuronavigation software (Brainsight, Rogue Research Inc, Montreal, Canada).

### 2.3 Data Processing

#### 2.3.1. COP Preprocessing

The center of pressure (COP) time-series data from the balance platform collected at 200 Hz were low-pass filtered (10 Hz), epoched into segment of 5 s including 1-s baseline rest period, mean-centered (zero-mean), and averaged across trials and conditions for each subject. The COP displacement was calculated as the trial average (in cm) of the cumulative distance traveled by the COP vector for the first 2 s of the perturbation in the forward/backward direction.

#### 2.3.2. EEG Preprocessing

The recorded EEG was processed offline using EEGLAB and BrainStorm toolboxes. First, we preprocessed the raw continuous EEG by down-sampling it into 250 Hz, followed by band-pass filtering between 1Hz and 50Hz using a Butterworth filter (4th order) to include all the frequency bands of interest. The electrical line noise of 60 Hz was removed using the Cleanline plugin of EEGLAB. Thereafter, the artifact subspace reconstruction (ASR) approach was implemented to correct the noisy bursts in the bandpass filtered continuous EEG data using the `clean_rawdata` plugin for EEGLAB [26]). The ASR approach contains bad channel rejection and noisy time window correction steps. The channels were removed if their correlation was less than 0.8 with the neighboring channels, and the burst detection criterion threshold (k in the ASR algorithm) was set at 20 based on the empirical study [27]. The artifactual time windows were removed if the fraction of total contaminated channels was above 0.25. After applying the common-average referencing to the ASR-corrected data, we performed the Independent Component Analysis (ICA) (extended Infomax algorithm)[28]. The resulting independent components were then classified into one of the seven categories ((1) Brain, (2) Muscle, (3) Eye, (4) Line noise, (5) Channel noise, (6) Heart, or (7) ‘other’) using a machine-learning tool ICLabel plugin of EEGLAB [29]. The dipole fitting was done using the DIPFIT tool in EEGLAB, which can be used to assess whether the dipoles corresponding to the ICs are ‘brain’-related or not.

We retained only those ICs that are classified as ‘brain’ (posterior probability *>* 0.5) and the residual variance (relative to the scalp topography) less than 20%. The selected ICs were back-projected onto the sensor space, which was used for EEG source localization.

#### 2.3.3. EEG Source Localization (ESL)

Since the brain activity recorded from the scalp gets attenuated due to the volume conduction effect (propagation of electric current flow through different layers of the brain and skull), it is recommended to estimate the cortical source activity [30]. To achieve this, we used the OpenMEEG tool [31] in the Brainstorm toolbox [32] to compute the forward head model, wherein a realistic head model comprising 4 layers (brain, inner skull, outer skull, and scalp surface) was reconstructed using individual T1 MRI scans and 3D EEG electrode positions using Brainsight Neuronavigation System (Rogue Research, Montreal, Canada). The volume conduction effect was simulated using the boundary element model (BEM) [33]. Once the forward model is obtained, we used the sLORETA [34] algorithm as a distributed source model to solve the ill-posed inverse problem. Estimating the solution using sLORETA requires the computation of the noise covariance matrix and the forward head model. Therefore, the noise covariance was estimated using the ‘baseline’ EEG, which corresponds to the pre-perturbation period (-1s to 0 s). After estimating the source-localized EEG, we parcellated the cortical surface into 68 anatomical regions using the Desikan-Killiany Atlas in the Brainstorm toolbox. Thereafter, the ROIs for the functional connectivity estimation and partial least squares analysis were determined based on our preliminary study [35] in which we identified the cortical regions with significantly more activation during the perturbation compared to the baseline period. To avoid bias concerning the hemispheric dominance, we included bilateral ROIs from the middle frontal gyrus, superior frontal gyrus, middle temporal gyrus, paracentral lobule, precentral gyrus, and postcentral, precuneus, cingulate gyrus, superior parietal lobule, and middle occipital gyrus.

#### 2.3.4. Clustering approach for stratification of TBI participants

To identify a neural marker of balance impairment within the TBI population, we dichotomized the TBI group into balance-impaired (BI-TBI) and balance-non-impaired TBI (BN-TBI) based on the COP displacement during the postural control task. Since there is no reference for COP-based stratification, we used the data-driven approach of unsupervised binary-class clustering using a one-dimensional k-means clustering algorithm. To avoid the bias of the choice of clustering algorithm, we also repeated the analysis with hierarchical agglomerative clustering algorithm with the ‘complete’ linkage criterion. In addition, we presented the findings of the stratification of TBI determined based on the BBS score since it is a well-known outcome measure of static balance with very high test-retest reliability [36].

#### 2.3.5. Functional Connectivity Estimation

The brain functional connectivity between two ROIs was calculated for the time segments (0-2 s) corresponding to the perturbation task and baseline period using the imaginary coherence (iCOH) [37]. Here, the time instant at which perturbation occurs is noted as *t* =0 s. The iCOH is considered a robust estimate of phase synchronization between two time-series data. To ensure computational tractability, we measured the iCOH between the “seed” voxels of different ROIs instead of every pair of voxels in an ROI. Formally the iCOH refers to the imaginary part of coherence, i.e., 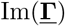, where, 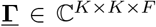 is a three-way tensor, where *K* and *F* are the number of ROIs and frequency bins of interest, respectively. Since the iCOH values are computed for each frequency bin, we averaged the iCOH values corresponding to the frequency bins within each frequency band (*θ* = 4 −8 Hz, *α* = 8 −13 Hz, *β* = 13 − 30 Hz). We also obtained the weighted node degree (WND or functional connectivity strength) for a given ROI by summing the 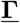 values of all its connections. In the context of graph theory, WND is one of the simplest and the most intuitive local network measures to evaluate the contribution of an ROI (or node) to the connectivity network [38].

### 2.4 Statistical Analysis using Mean-centered Partial Least Squares Correlation (MC-PLSC)

Partial Least Squares Correlation (PLSC) for neuroimaging applications was introduced by McIntosh [39]. The PLSC algorithm is a multivariate technique that is used to explore the statistical association between two sets of variables. Depending on the research question, we can choose one of the variants of PLSC, such as behavior PLSC (brain- and behavior measures as two different variables), task PLSC (brain measures corresponding to two different conditions/tasks such as attention vs. rest), or seed PLSC (to analyze the functional connectivity in a specific brain region, regarded as the seed) [23]. The PLS algorithm has the inherent advantage of dealing with more variables (*P*) than the number of observations (or samples *N*), which is a befitting choice for most neuroimaging studies. In this study, we used the Mean-Centered task PLSC (MC-PLSC), which is conceptually very similar to Barycentric Discriminant Analysis (BADA) [40] in the sense that MC-PLSC allows us to find a set of brain features that best maximizes the absolute differences between the groups. In the MC-PLSC, the group labels are used as dummy-coded categorical variables. For a thorough treatment of various types of PLSC, we direct the readers to a review article by [23].

The mean centered PLSC procedure followed in this study is briefly explained below.

A brain measure matrix **X** ∈ ℝ^*N ×P*^ is built by concatenating the rows of brain imaging features from *N* subjects with *P* being the total number of features considered for a subject. In the primary analysis, *P* = 20, because *P* denotes the summed functional connectivity strength of 20 ROIs, which are computed from the functional connectivity matrix constructed with the EEG data of each participant. The binary-valued dummy coding matrix is denoted as 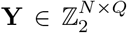, where *Q* is the number of experimental groups (in our case, *Q* = 3). After Z-score normalization of **X**, its group-wise average is computed as **M** = diag(**1**^⊤^**Y**)^−**1**^**Y**^⊤^**X**, where **1** is the vector of 1s of length *N*, and ⊤ is the matrix transposition operator. This matrix scales the X according to the mass of **X** (the values are scaled such that the sum of the masses is equal to one). Thereafter, the mean-centered matrix (also called cross-block covariance matrix) is computed as 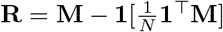, where **1** is a N-length vector of 1s [23].

The singular value decomposition (SVD) of **R** is expressed as **R** = **UΔV**^⊤^ where **U** and **V** are left and right singular column vectors (**u**_*k*_ and **v**_*k*_, *k* ∈ {1, 2, …, *Q*}, respectively, and the diagonal matrix **Δ** contains the singular values *δ*_*k*_. In the PLSC literature, the singular vectors **U** and **V** are termed design/group salience and brain salience, as they reflect the group contrast and weighted contribution of brain features, respectively. The projection of **Y** onto the design salience **U** is known as the design scores, **L**_*Y*_ = **YU**. Likewise, the projection of **X** onto the brain salience **V** results in the brain scores, **L**_*X*_ = **XV**. The mean centered PLSC effectively decomposes **R** into *k* components that optimally separate the groups by finding pairs of group/design and brain latent vectors with maximal covariance.

We performed nonparametric permutation testing to assess the statistical significance of LVs from the PLSC analysis. In this procedure, the brain imaging features are randomly shuffled many times (1000 in our case), and the LVs are computed using the PLSC every time, thus disregarding the original relationship between brain imaging (**X**) and group label data (**Y**). The latent variable *k* is considered significant if the empirical singular value (*δ*_*k*_) is higher than 95% of the values obtained from the null distribution. Provided the LV is significant, we tested which of the loading weights () are robust amongst the brain connectivity features. For this, we conducted a bootstrap procedure wherein the rows of the X and Y were sampled with replacement. The saliences with the standard error, 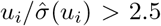 and 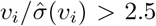, from bootstrap samples (1000 in our study) are deemed significantly stable. Since the permutation and bootstrapping procedure can affect the alignment (rotation) of the LVs, we applied Procrustes rotation to the LVs so that they correspond to the original data [41]. The MATLAB scripts related to this study were adapted from the myPLS toolbox: https://github.com/danizoeller/myPLS.

#### Confidence Intervals

To test whether the separation between groups was significant, we used generalized Principal Component Analysis to project the factor scores [40]. The factor scores given by *F* = **UΔ** were projected onto a 2-dimensional map with the Principal Component Analysis. A 95% confidence interval of these factor scores was obtained using bootstrapped sampling, where the factor scores were obtained for every sampling.

### 2.7. Analysis of Variance

To determine if there were statistically significant differences in the network functional connectivity strength across the three groups, we conducted a one-way analysis of variance (ANOVA) for each network, i.e., Default Mode Network (DMN), Frontoparietal Network (FPN), Sensorimotor Network (SMN), and Visual Network (VN). The group was considered a between-subject factor in each analysis, comprising three levels: BI-TBI, BN-TBI, and HC. The dependent variable was network strength, representing the strength of within-network functional connectivity, which was computed as the weighted sum of connectivity values between regions belonging to the same network. The assumption of homogeneity of variances was tested using Levene’s test. If significant differences were found in ANOVA results, post-hoc comparisons using Tukey’s Honest Significant Difference (HSD) test were conducted to identify the groups that differed significantly.

## 3 Results

### 3.1 Univariate Analysis of Balance Functional Measures

In this section, we compare behavioral measures based on BBS and COP displacement measured during high backward perturbation. While the COP displacement measures the ability to maintain balance in response to balance perturbation dynamically, BBS is a measure of static and dynamic functional balance and thus provides complementary information [42]. A univariate analysis based on a two-tailed t-test revealed that the individuals with TBI (mean *±* SD = 11.64 *±* 4.28, 95% CI =[4.79, 19.18]), as a whole group, showed significantly larger COP displacement (*t* = 3.07, *p* = 0.004, Cohen’s D = 1.09) than HC (mean *±* SD = 7.82 *±* 2.33, 95% CI = [4.7, 13.13]). The univariate analysis of BBS data was done using the nonparametric Wilcoxon Ranksum test, given the non-normal distribution, which showed that the HC participants have a significantly higher BBS scores (median = 56, range = [55, 56]) than the TBI (median, range = 51, [34, 56]), with *p* = 0.007 and z = 2.68.

### 3.2 Clustering-based Dichotomization of TBI

To characterize the level of balance deficits across TBI participants based on their COP displacement values, we used an unsupervised clustering approach using 1-D k-means clustering described in **Section 2.3.4**. The scatterplot of COP displacement values and BBS scores of all the participants in both groups are shown in **Fig. 2**(a). The scatterplot to illustrate the clustering results of COP displacement values is shown in **Fig. 2**(b), where the one-dimensional data of COP displacement values from all 17 TBI individuals are clustered into either balance impaired (BI-TBI_COP_) or balance non-impaired (BN-TBI_COP_) TBIs. The cluster threshold was determined as the midpoint between the centroids of the two clusters (threshold value of 11.9 cm), we obtained the subgroup sample of BN-TBI_COP_ (*N* = 9) and BI-TBI_COP_ (*N* = 8).

**Figure 2:**
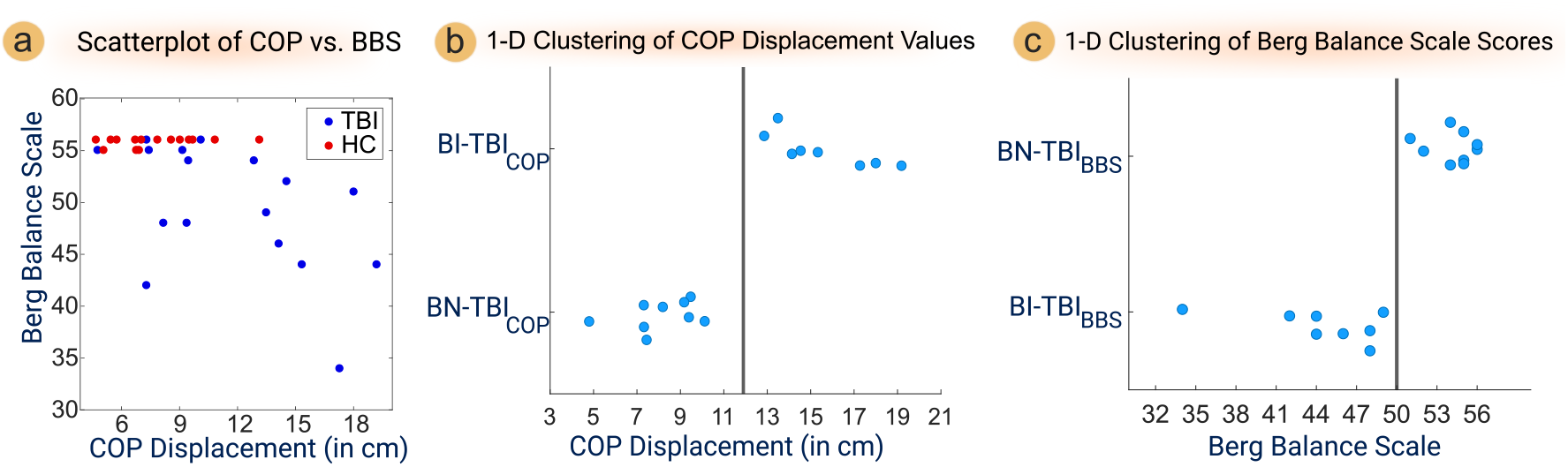
**(a)** Scatterplot of COP displacement (x-axis) and BBS scores (y-axis) from TBI and HC groups. **(b)** The scatterplot of COP data was clustered into two groups (BI-TBI_COP_ and BN-TBI_COP_) with the *k* -means method. **(c)** The scatterplot of BSS data was clustered into BI-TBI_BBS_ and BN-TBI_BBS_ groups using the same approach. The vertical line indicates the cluster threshold to assign the TBI individuals into one of the two categories.

Similarly, we also determined the cluster threshold for the BBS score-based clustering as 49, as it is the midpoint between two cluster centroids (**Fig. 2**(c)), in line with Shumway-Cook et al. (1997) that suggested a BBS score of 49 as the cutoff score to determine individuals with vs. without risk of falls. Equivalently, we designated the individuals with and without risk of falls as BI-TBI_BBS_ (*N* = 8) and BN-TBI_BBS_ (*N* = 9), respectively.

We conducted a one-way ANOVA to assess whether there were any group differences in age, given its role in balance and postural control. When we compared the COP-stratified groups, one-way ANOVA revealed no significant differences in the age across any groups (F (2, 29) = 1.48, *p* = 0.2443). Similar analysis for the BBS-stratified groups also resulted in the same conclusion (F (2, 29) = 0.07, *p* = 0.9313).

### 3.3 Group-level Comparison of the Postural Task-relevant Functional Connectivity

To study the effect of groups on the theta-band functional connectivity pattern between different ROIs, we compared the within-network connectivity strength across groups. Levene’s test for unequal variance between the groups for each network showed no significant differences in the variance (DMN: *p* = 0.504, Levene’s statistic = 0.702; FPN: *p* = 0.2, Levene’s statistic = 1.705; SMN: *p* = 0.752, Levene’s statistic = 0.288; VN: *p* = 0.27, Levene’s statistic = 1.348). The heatmap figure illustrating the theta-band is presented in **Fig. 3**(a), where the relatively weaker connectivity strength pattern can be seen in the BI-TBI_COP_ compared to the BN-TBI_COP_ and HC groups. The quantitative comparison with one-way ANOVA revealed a non-significant main effect of the group on the within-group DMN network strength differences (F (2,29) = 0.53, ***p*** = 0.595, *η*^2^ = 0.04). Similarly, for FPN and VN, the one-way ANOVA did not show a significant main effect of the group [F (2,29) = 1.82, *p* = 0.180, *η*^2^ = 0.11) and F (2,29) = 1.59, *p* = 0.222, *η*^2^ = 0.10, respectively]. However, the SMN showed a large effect of the group on the network strength [F (2,29) = 3.27, p = 0.052, *η*^2^ = 0.18]. The data showed a pattern of smaller network strength within the SMN in BI-TBI_COP_ compared to the HC, as demonstrated by the post-hoc t-test upon correction for multiple comparisons using Tukey’s HSD test (*p* = 0.047). The within-network connectivity strength results are presented as half-violin plots overlaid on the boxplot to illustrate the distribution of data points in **Fig. 3**(b).

**Figure 3:**
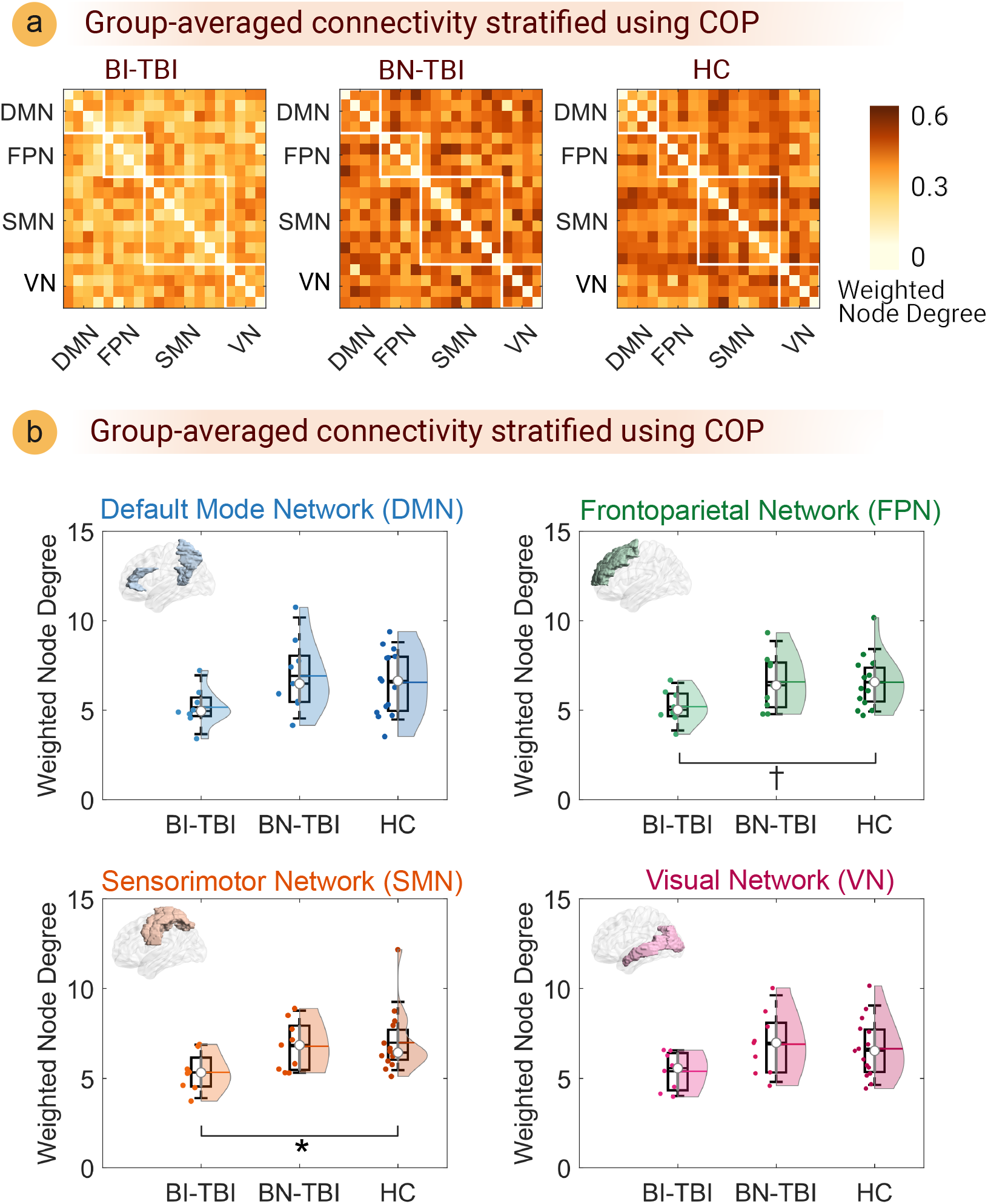
**(a)** Group-averaged theta-band functional connectivity patterns are shown as heatmaps. The groups are dichotomized into BI-TBI_COP_ and BN-TBI_COP_ based on the 1-D clustering of COP values using hierarchical agglomerative clustering. Each element on the connectivity matrix represents the group-averaged connectivity value between two cortical ROIs measured using the theta-band coherence. Different cortical ROIs are grouped into the following intrinsic brain functional networks: DMN, FPN, SMN, and VN. **(b)** Between-group comparison of the network strength (or the summed weighted node degrees) for each functional network represented by the cortical ROIs.

### 3.4 Mean-centered Partial Least Squares Correlation (MC-PLSC) Analysis

We present the findings from MC-PLSC analysis using theta-band functional connectivity strength of cortical ROIs and group stratification (BI-TBI, BN-TBI, and HC) based on the COP criterion. Per the permutation testing of singular values, only the first LV was significant [the empirical singular value exceeded 98.7% of the histogram distribution (p = 0.013), as shown in Fig. 4(a)], and it accounted for 91.8% of the cross-block covariance. The corresponding design saliences and their bootstrapped confidence ellipses are plotted in Fig. 4(b) and 4(c). A similar analysis using BBS stratification resulted in one significant LV (p = 0.04, per the permutation testing shown in Fig. 4(d)), which explains nearly 90.4% cross-block covariance.

**Figure 4:**
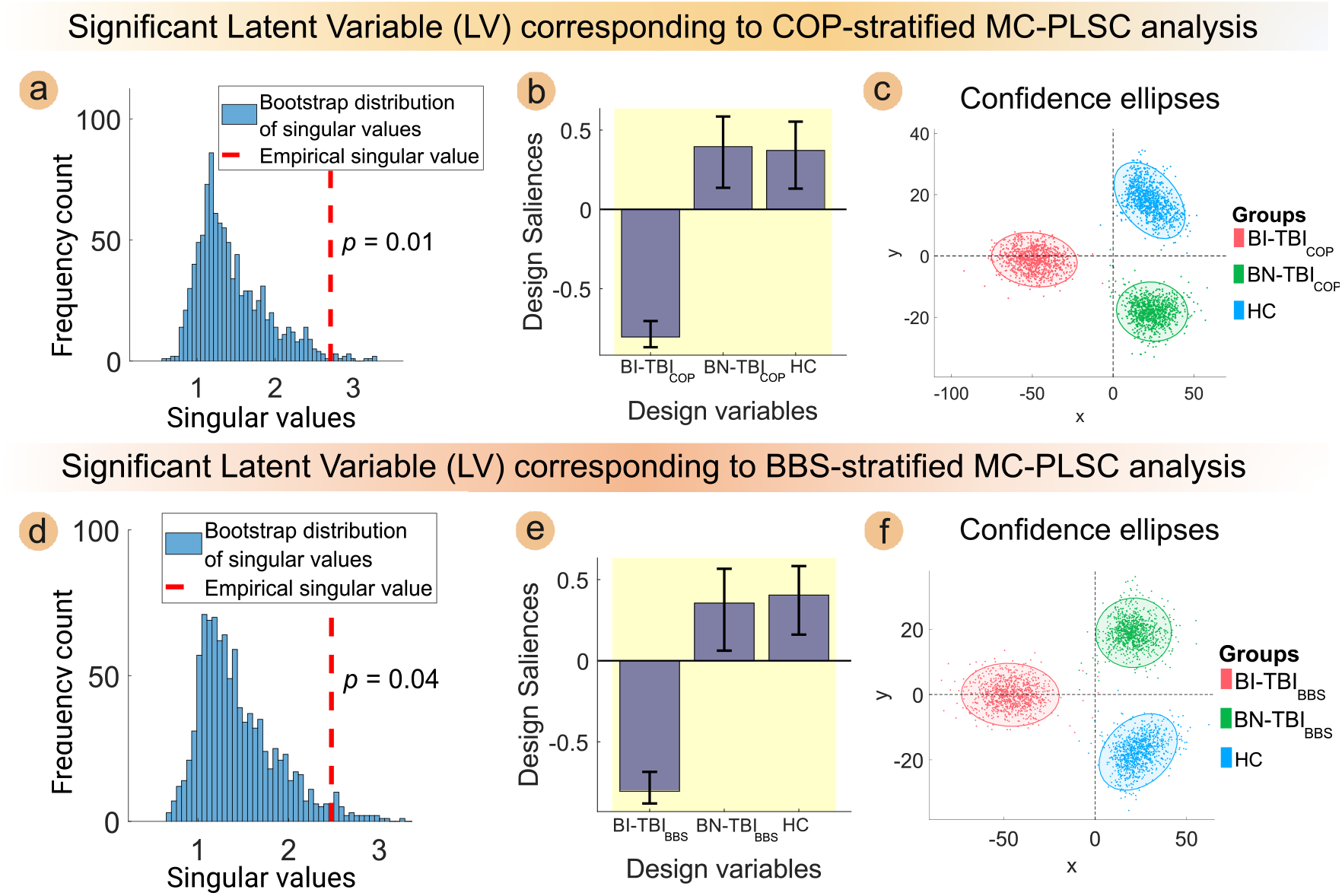
The results from the MC-PLSC, when implemented with the individuals stratified based on COP and BBS, are illustrated in the top and bottom rows, respectively. The brain imaging variable is the theta-band functional connectivity strength of each of the ROIs, and the design variable is the group label. In **(a)** and **(d)**, the empirical singular value associated with the first LV (red dotted line) is overlaid on the histogram of singular values (blue bar graph) computed from the permutation sample. The empirical singular value is deemed statistically significant in each instance since it remains above the 95th percentile of the respective null distribution. The design salience U is represented by a bar plot in **(b)** and **(e)**, where the height of each bar denotes the magnitude of an element of the first left singular vector. From **(b)** and **(e)**, one can infer that BI-TBI significantly differs from BN-TBI and HC. The error bars generated with the respective singular vectors from the bootstrap sample are shown along with the bar plots to mark the 95% confidence interval. In both cases, all the elements of design salience are considered robust by the bootstrap test. The 95% confidence ellipses of scatter plots in **(c)** and **(f)** are obtained with the first and second LVs. There is a clear separation between BI-TBI and the combined set of BN-TBI and HC along the x-axis (first LV), but not along the y-axis (second LV).

The MC-PLSC analysis of theta-band brain connectivity in COP-stratified and BBS-stratified individuals found one LV in each case. The LV effectively distinguishes the three groups (BI-TBI, BN-TBI, and HC) when contrasting BI-TBI with BN-TBI and HC combined [**Fig. 4**(b) and **Fig. 4**(c)]. This observation suggests that the theta-band neural response for BN-TBI resembles that of HC, despite the stratification strategy [**Fig. 4**(e) and (f)], thus supporting our hypothesis and rationale for separating TBIs into balance-impaired and non-impaired populations.

Upon bootstrap testing of the brain salience—the right singular vector corresponding to the singular value that survived the permutation test—we found five ROIs that are robustly associated with distinguishing the BI-TBIs from the rest, regardless of the type of stratification as shown in **Fig. 5**(a). The “robustness” of the brain imaging feature is evaluated based on the Bootstrap Ratio (BSR), i.e., BSR *>* 2.5, meaning that the salience values or weighted contributions are non-zero with 99% confidence interval. Specifically, we observed the theta-band brain connectivity between the following ROIs—*left middle frontal gyrus, right paracentral lobule, precuneus*, and *bilateral middle occipital gyri* —as “stable features” from both the COP-stratified and BBS-stratified analysis.

**Figure 5:**
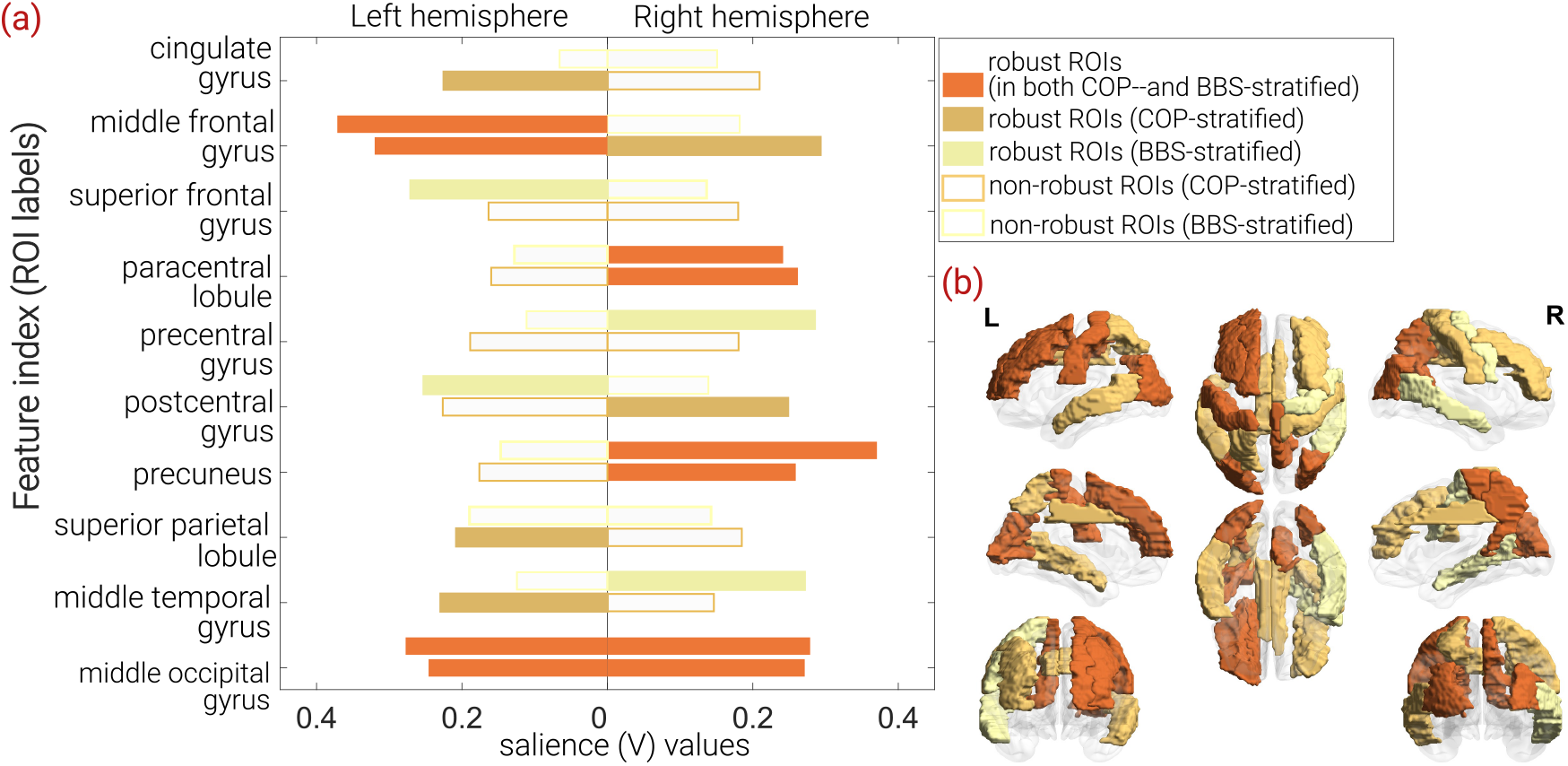
(a) The bar plot of brain saliences (right singular vector v 1 resulting from the SVD of R) associated with the significant LV. The top bar for each ROI denotes the magnitude of the brain salience for BBS-stratified analysis, whereas the bottom one corresponds to the COP-stratified analysis. The brain saliences were selected by the bootstrap test based on the criterion that BSR must exceed 2.5. (b) The diagrammatic representation of ROIs germane to stable theta-band brain connectivity overlaid on the cortical surface. (b) Corresponding visualization of robust ROIs overlaid on the cortical surface. The figure was generated using the BrainNet Viewer MATLAB tool [43].

Similar analyses were conducted for alpha- and beta-band connectivity features as well. However, the MC-PLSC did not reveal any significant LV that disentangles the three groups. These results are presented in the supplementary material.

## 4. Discussion

### 4.1. Disentangling the Levels of Balance Impairment in TBIs

Stratifying individuals with TBI into balance-impaired and non-impaired groups helps better understand the clinical relevance and prognostic significance of balance deficits. For identifying people with low vs. high risk of falls, an unsupervised clustering approach using the lower-extremity Fugl-Meyer Assessment score [44] and a supervised classification approach using the BBS score [45] have been studied. Our analysis relies on classifying the COP displacement that measures the task-specific postural sway with an unsupervised clustering, and the results showed that the individuals with TBI can reliably be stratified into two groups. Contrary to our expectation, assigning subjects into two groups did not remain the same when the stratification criterion was COP and BBS. The reason could be that the BBS is a subjective clinical assessment scale that strongly correlates with the gross functional outcome measure (e.g., Timed Up-and-Go) rather than the laboratory measures of body sway (COP displacement), which can be sensitive to platform perturbation parameters. Therefore, we cannot risk conflating these two disparate measures to jointly evaluate the group stratification using a 2-D clustering approach. Moreover, there may not be clearly defined clusters in the 2-D space of these two variables, and therefore, stratifying participants is better suited to one variable at a time. In our study, *k* -means clustering of COP displacement, a threshold of 11.9 cm, would separate the TBI participants into two groups—balance impaired and balance non-impaired. A similar clustering-based approach to stratify the participants using their BBS scores resulted in a threshold of 49. This threshold is in line with the findings of [45] and [46], who identified people with high fall risk and physically inactive participants, respectively. Note that since our experimental task uses study-specific parameters relevant to the backward perturbation of the Neurocom platform, the threshold for the COP displacement (i.e., 11.9 cm) for group stratification may not be generalized. Future studies must consider COP-based metrics that are robust to the choice of posturography platforms.

### 4.2 Role of Different Cortical Regions in the Neurophysiological Control of Balance

Regarding the neural underpinnings of balance impairment in individuals with chronic TBI, recent studies suggest that such complaints are predominantly due to central sensory system dysfunctions rather than issues within the peripheral vestibular or oculomotor systems [47]. The subsequent sections delve into the involvement of various brain regions in the mechanisms of postural control in TBI.

To encapsulate the findings within the framework of the closed-loop mechanism of balance control, we reference the hypothetical model by Takakusaki [15]. This model introduces the “body schema,” illustrating the participation of multiple brain regions in balance control. According to Takakusaki’s posture-gait control model, sensory signals from the visual cortex, vestibular cortex, and primary sensory cortex are received by the midbrain and subcortical regions, including the cerebellum, brainstem, thalamus, and cerebral cortex. These signals are then processed by the temporoparietal cortex to construct the body schema. Specifically, the temporoparietal cortex facilitates the generation of motor control commands from the supplementary motor area (SMA) and premotor (PM) regions, aided by the basal ganglia and cerebellum. In our study, the middle frontal gyri, encompassing the SMA/PM region, demonstrate a notable role in the BI-TBI group compared to the BN-TBI and HC groups, as indicated by its significant BSR in the MC-PLSC model derived from theta-band WND features. The functional significance of the middle frontal gyri in anticipatory postural control is underscored in previous research [48], and their involvement in the supraspinal motor network for stance and locomotion in elderly adults is documented [49].

Following the reception and processing of sensory signals by the visual, vestibular, and somatosensory cortices, the paracentral lobule and precentral gyrus manage the motor commands for balance control, corresponding to the leg region of the primary motor cortex (M1) [50]. Previous studies have elucidated the role of the paracentral lobule as a region that processes the motor commands for balance control after receiving the sensory inputs from the visual, vestibular, and somatosensory cortices [51]. In the context of theta-band functional connectivity, the SMA located just in front of the paracentral lobule is associated with pull perturbation while standing [52]. Additionally, the paracentral lobule and precuneus are pivotal in interconnecting various brain regions due to their anatomical links, facilitating efficient pathways for integrating different brain functions. Notably, the reduced cortical activity in precuneus may be associated with a misperception of motion in individuals with persistent postural-perceptual dizziness [53], a condition affecting one in three TBI survivors[54].

Within the context of the posture-gait control model in [15], the superior parietal region is implicated in anticipatory postural adjustments by detecting postural instability [55] . Furthermore, theta-band coherence-based connectivity analysis reveals significant functional connections with the left superior parietal lobule, alongside the right paracentral lobule and left postcentral gyrus. Given the roles of the cingulate and angular gyrus in dynamically regulating attention to unpredictable events [56], and their anatomical connections to the basal ganglia and cerebellum, it is anticipated that postural control signals from motor regions are relayed to the cortico-reticular and reticulospinal tracts via the cingulate gyrus.

In terms of visual perception related to balance perturbation, the occipital lobe regions are presumed to play a crucial role in discerning static versus dynamic motions (or tilts) of the posturography platform [57]. Aligning with this discussion, the middle occipital gyrus, as identified by theta-band WND features in our study, is expected to significantly contribute to the sensory integration of visual and motor functions [58]. Additionally, activation of the middle temporal gyri has been observed in simulations (or imaginations) of postural control tasks [56]; [59], with the activated areas closely related to the parietal insular vestibular cortex, a region generally acknowledged for processing vestibular signals pertinent to postural control [15].

### 4.3 Roles of the Functional Networks Identified by PLSC

For qualitative assessment of our findings, we now discuss the roles of functional networks to which the aforementioned ROIs belong. Although it is not trivial to assign the anatomical ROIs to specific functional networks, a recent study has tested the spatial correspondence between the anatomical regions (based on the Desikan-Killiany atlas) and functional networks (based on the Yeo-Atlas) [60]. On this premise, we explored the role of the significant ROIs returned by the MC-PLSC analysis in the context of functional networks involved in postural control. In this regard, our focus is mainly on the two intrinsic functional networks: SMN and VN, as these networks showed high spatial correspondence with the motor and visual areas per the Desikan-Killiany atlas [60]. Not surprisingly, the SMN is reported to play the most critical role during postural control mechanisms by facilitating sensorimotor integration [61]. While the roles of different regions within the SMN are widely studied in postural control [62]; [50]; [48], the network-level mechanisms are not well studied in the TBI population. Our study offers preliminary evidence that the functional connectivity network strength of SMN comprising bilateral paracentral lobule is capable of discriminating the BI-TBI from BN-TBI and HC. Moreover, the middle occipital gyrus (MOG) has also been found to be a robust brain salience in our study, presumably because the MOG is involved in the visuospatial perception to regain balance control in response to the platform perturbation [63]; [64]. We also noticed that the left superior parietal lobule (SPL) contributes to distinguishing the groups based on the COP displacement. The role of left SPL in motor attention tasks is highlighted in [65]. The SPL is reported to be preactivated during the anticipatory movement which will subsequently predict performance to upcoming targets [66].

### 4.4 Role of Frequency Bands

Our main findings of the MC-PLSC approach to disentangle balance impairment levels are related to the theta-band connectivity features. Some of the concordant evidence demonstrating the role of the theta band in balance tasks include [67]; [52]; [68]. On the other hand, we did not find any significant LV pertaining to alpha or beta frequency bands. This observation contradicts the general notion that alpha and beta frequency bands are associated with balance tasks. For example, the average activity in alpha and beta frequencies was significantly modulated during dual-task walking conditions, indicating an increased cognitive load during walking [69]. In addition, task-dependent alterations of the alpha frequency band during a motor interference task were reported in [70], where participants had to walk while balancing a stick with interlocked rings at the end. This finding suggests that the alpha-band activity is involved in cognitive-motor interference, which is a crucial aspect of postural control during complex motor tasks. Regarding the role of the beta band, the greater beta suppression is shown to reflect less postural sway (or better balance ability) [71]; [72]. However, most of these studies have only analyzed the variations in the band-specific spectral power in healthy volunteers and not the corticocortical functional connectivity, especially in neurological populations. The most relevant literature evidence regarding the EEG connectivity during balance control tasks is found in [73]; [17], where it was reported that the increased balancing difficulty resulted in a significant increase in connectivity strength in theta, alpha, and beta bands. On the contrary, in our study, the increased level of balance impairment is shown to reflect poor functional connectivity strength (**Fig. 3**). One explanation for this discordant observation is that the studies [73]; [17] have a within-group analysis of varying difficulty levels, but ours is the between-group analysis of the same difficulty level. We recommend that future studies comprehensively analyze the functional connectivity differences between balance-impaired- and non-impaired populations for different levels of task difficulty to reach a conclusion.

### 4.5. Limitations and Future Directions

We acknowledge the limitations of our study that warrant further investigation. First, our sample size is relatively small for a stratified analysis but reasonable in terms of the sample size presented in [22]. Second, the COP displacement threshold for the group stratification is specific to our study settings, which needs to be ascertained by a much larger sample. Third, the future study design of postural control and gait-related experimental tasks in TBI must consider the level of impairment using a more sensitive balance functional measure, which is robust to ceiling effects (e.g., Mini Balance Evaluation Systems Test and Functional Gait Assessment) and not based on just the level of severity (mild/moderate/severe) assessed by the Glasgow Coma Scale at the time of injury. Fourth, regarding the ROIs, we did not study the task-specific activity of deep sub-cortical neural substrates such as the brainstem, basal ganglia, pedunculopontine nucleus, and cerebellum, which are also shown to be involved in the postural control [51]. Finally, given the limited accuracy of EEG source localization of subcortical structures, we recommend further research in this direction.

We posit that the knowledge of the neural correlates of balance deficits can be applied to advance the field of neurorehabilitation by targeting the key brain regions with specific interventions, e.g., neuromodulation treatments targeting the leg motor cortical regions and neurofeedback games engaging the sensorimotor regions. For instance, cerebellar intermittent theta burst stimulation is shown to be a promising intervention to improve gait and balance in individuals with stroke [74].

## Conclusion

In this study, we have presented for the first time a stratified analysis of balance deficits in TBI using brain connectivity features pertaining to a balance perturbation task. Since the heterogeneity in TBI poses challenges in identifying robust brain imaging features correlated with the impairment, we employed a multivariate statistical framework, namely MC-PLSC. Importantly, when the theta-band functional connectivity strength of specific brain ROIs were used as the brain imaging features in MC-PLSC, the algorithm could reliably distinguish balance-impaired TBIs from balance-non-impaired TBIs and healthy controls. Our interpretation is that the selected regions, namely, *left middle frontal gyrus, right paracentral lobule, precuneus*, and *bilateral middle occipital gyri*, play a critical role in postural control, and their weakened functional connectivity during the postural control task may reflect maladaptive balance performance. Understanding the role of key ROIs may help design novel therapeutic interventions (e.g., neuromodulation and/or goal-directed movement therapies) for improving the balance functions in TBI.

## ACKNOWLEDGMENT

This study was supported by funding from the multi-investigator research grant awarded by the New Jersey Commission on Brain Injury Research (grant No. CBIR15MIG004). In addition, we also acknowledge the funding support from the Advanced Rehabilitation Research and Training (ARRT) program funded by the National Institute of Disabilities, Independent Living, and Rehabilitation Research (grant No. 90ARHF0002). Furthermore, we thank our research Engineer, Armand Hoxha, for assisting with the study data collection.

## DATA AVAILABILITY STATEMENT

The raw data involving the patient information is not publicly available due to the IRB restrictions. However, the MATLAB scripts and the deidentified preprocessed EEG and COP data supporting this study will be made available from the corresponding author upon reasonable request.

## CRediT AUTHORSHIP STATEMENT

The following color-coded table illustrates the contributions of each author based on the CRediT taxonomy (Brand *et al* 2015). The level of contribution is colored coded in terms of ‘support’ (light blue), ‘equal effort’ (blue), and ‘lead’ (dark blue).

