## Supplementary Material for "Identifying Neural Correlates of Balance Deficits in Traumatic Brain Injury Using Partial Least Squares Correlation Analysis"

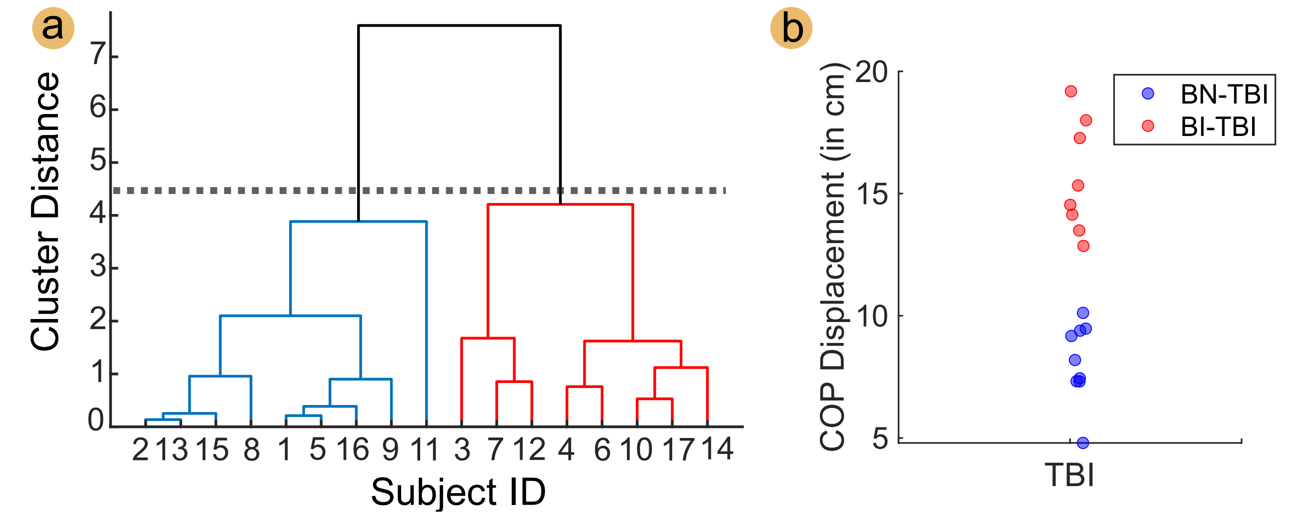


**Figure S1**: (a) Hierarchical agglomerative clustering (HAC) of the COP displacement values to stratify the TBI participants into two groups, color-coded in blue and red. The x- and y-axis denote the subject ID and the distance between clusters, respectively. The distance threshold between the cluster centroids to return two clusters is marked with the black dotted line. The tree branches in red and blue denote the balance impaired TBIs (BI-TBI) and balance non-impaired TBI (BN-TBI), respectively. We note that the cluster assignment for each subject was the same for both methods (*k*-means and HAC). (b) The color-coded scatterplot shows the COP displacement values of BI-TBI (red) and BN-TBI (blue).

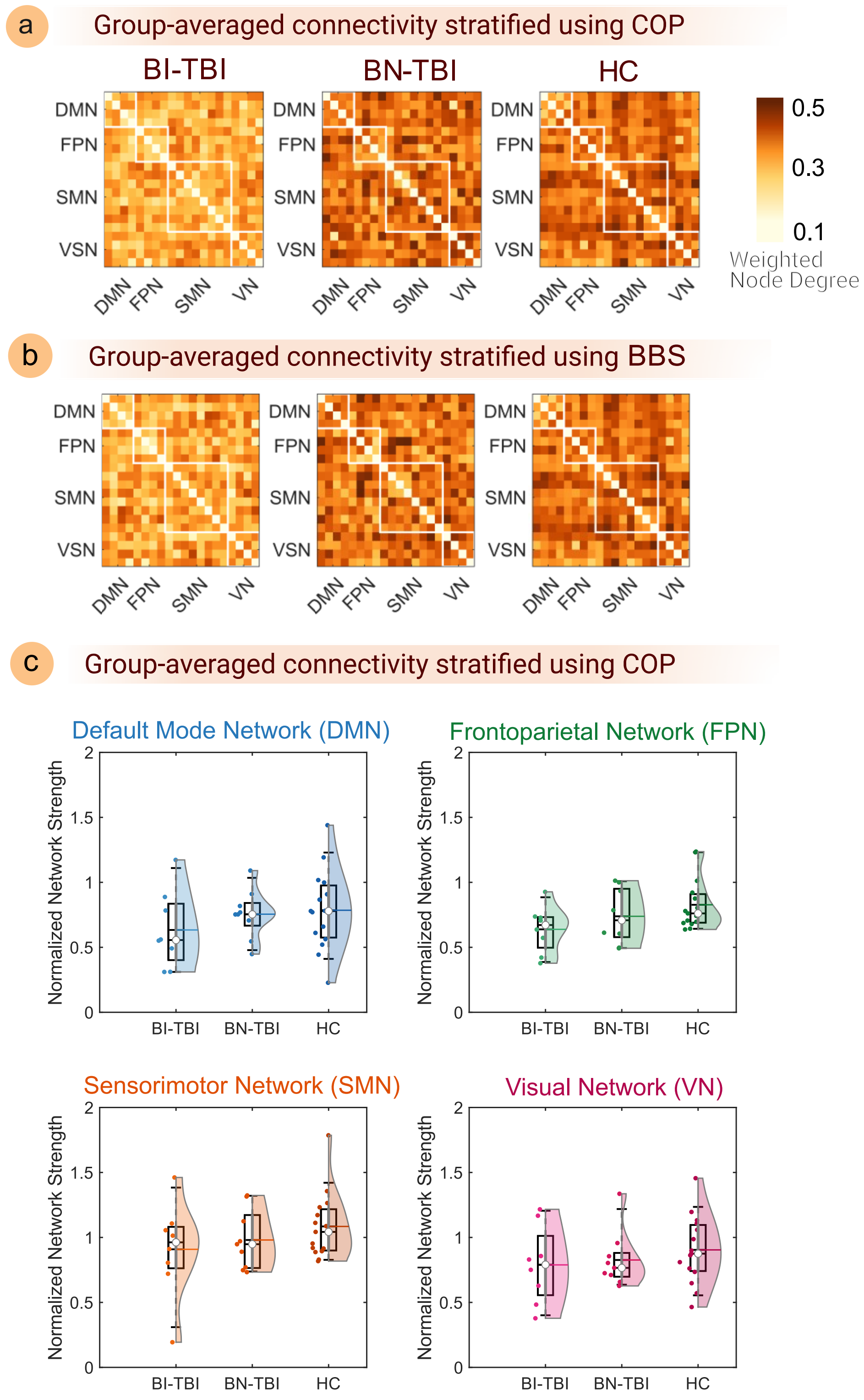


**Figure S2**: Group-averaged **alpha**-band functional connectivity patterns are shown as heatmaps for (a) the HC and COP-stratified TBI participants, and (b) HC and BBS-stratified TBI participants. Each element on the connectivity matrix represents the group-averaged connectivity value between two cortical ROIs measured using the **alpha**-band imaginary part of coherence. Different cortical ROIs are grouped into the following intrinsic brain functional networks: DMN, FPN, SMN, and VN. (c) Between-group comparison of the network strength (or the summed weighted node degree) for each functional network represented by the cortical ROIs. No statistically significant differences were found between the groups after correcting for multiple comparisons using Tukey's HSD test following the one-way ANOVA for any network (see **Table S1** for statistical results of the one-way ANOVA).

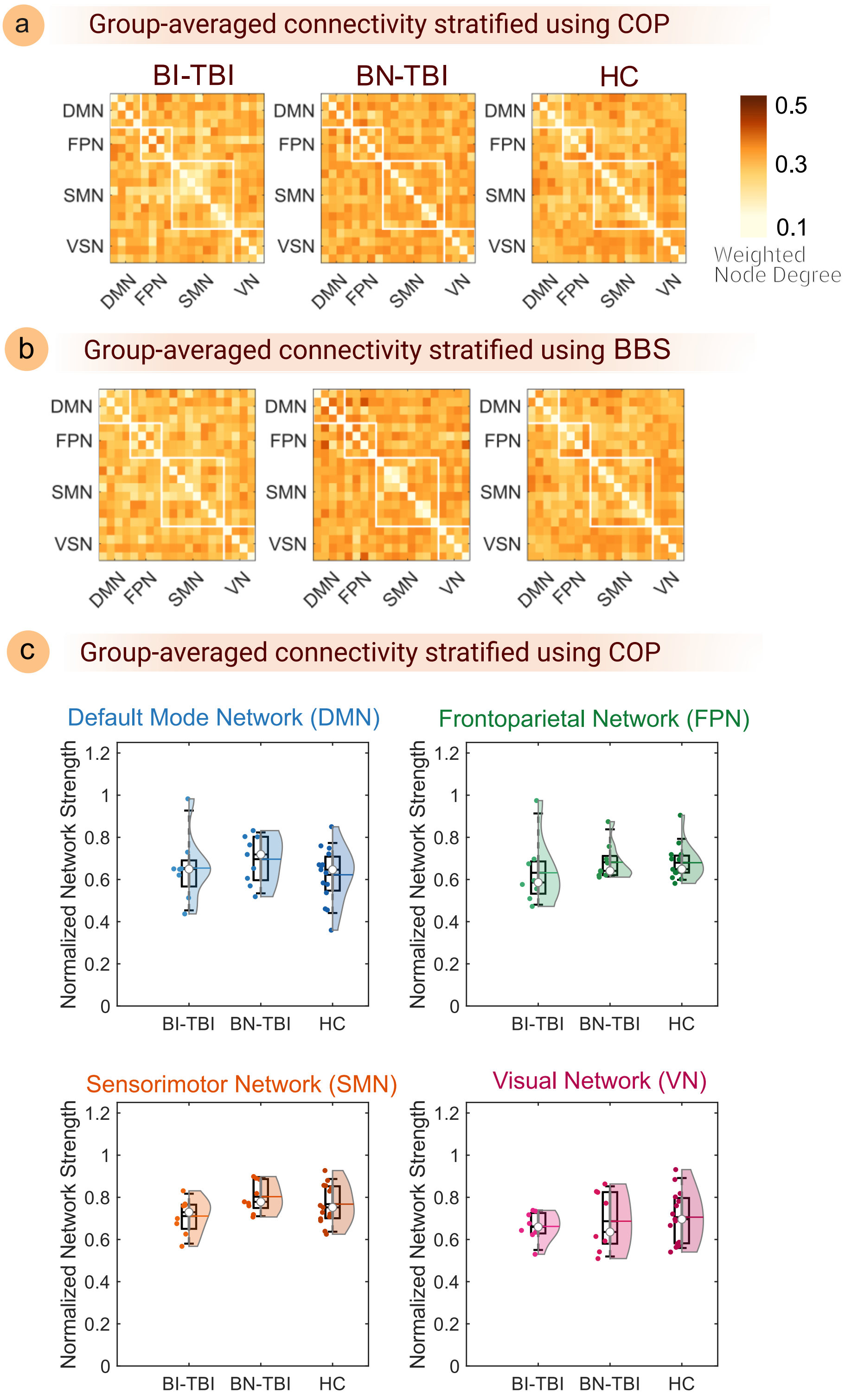


**Figure S3**: Group-averaged **beta**-band functional connectivity patterns are shown as heatmaps for (a) the HC and COP-stratified TBI participants, and (b) HC and BBS-stratified TBI participants. Each element on the connectivity matrix represents the group-averaged connectivity value between two cortical ROIs measured using the **beta**-band imaginary part of coherence. Different cortical ROIs are grouped into the following intrinsic brain functional networks: DMN, FPN, SMN, and VN. **(c)** Between-group comparison of the network strength (or the summed weighted node degree) for each functional network represented by the cortical ROIs. No statistically significant differences were found between the groups after correcting for multiple comparisons using Tukey's HSD test following the one-way ANOVA for any network (see **Table S1** for statistical results of the one-way ANOVA).

**Table S1: Summary of one-way repeated measures ANOVA to study the main effect of group on the alpha and beta-band network strength as dependent variables.**

| Brain functional networks of interest | Frequency bands of interest | Group main effect | | |
|---|---|---|---|---|
| | | F (2,29) | p-values | Effect size (partial $\eta^2$) |
| DMN | Alpha | 0.81 | 0.455 | 0.05 |
| | Beta | 0.86 | 0.435 | 0.06 |
| FPN | Alpha | 2.51 | **0.098** | 0.15 |
| | Beta | 0.64 | 0.538 | 0.04 |
| SMN | Alpha | 1.11 | 0.343 | 0.07 |
| | Beta | 2.46 | **0.103** | 0.15 |
| VN | Alpha | 0.59 | 0.562 | 0.04 |
| | Beta | 0.38 | 0.690 | 0.03 |

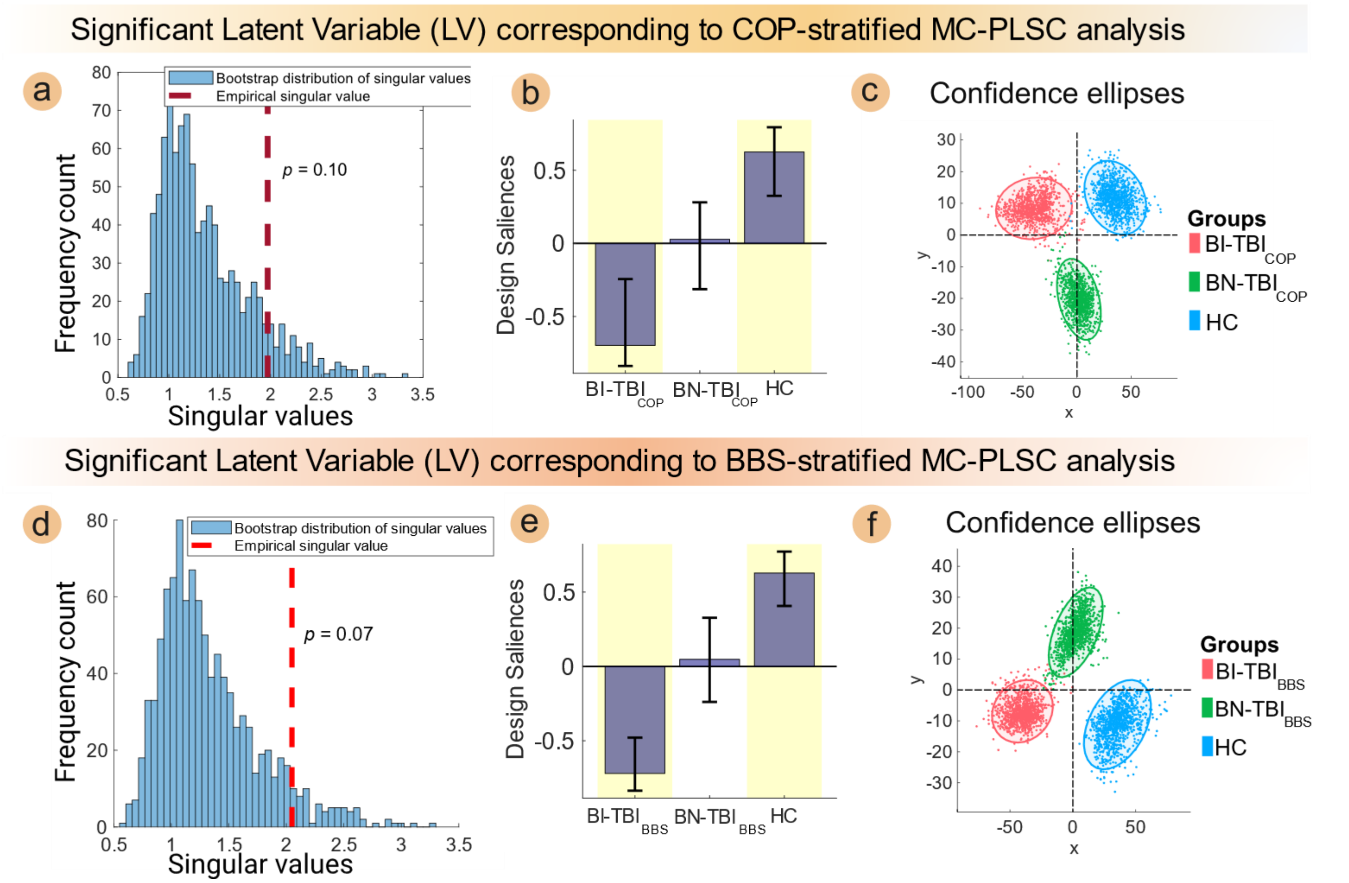


**Figure S4**: The results from the MC-PLSC, when implemented with the individuals stratified based on COP and BBS, are illustrated in the top and bottom row, respectively. The brain imaging variable is the **alpha**-band weighted node degree, and the design variable is the group label. In (a) and (d), the empirical singular value associated with the first LV (red dotted line) is overlaid on the histogram of singular values (blue bar graph) computed from the permutation sample. In each instance, the empirical singular value is deemed statistically not significant since it remains below 95th percentile of the respective null distribution. The design salience **U** is represented by a bar plot in (b) and (e), where the height of each bar denotes the magnitude of an element of the first left singular vector. From (b) and (e), one can infer that BN-TBI is not significantly different from either BN-TBI and HC, as denoted by the white background. The error bars generated with the respective singular vectors from the bootstrap sample are superimposed on the bar plots to mark the 95% confidence interval. In both cases, BI-TBI and HC design saliences were robust (yellow background) as their bootstrap ratio was more than 2.5 unlike BN-TBI. The 95% confidence ellipses of scatter plots in (c) and (f) are obtained with the first and second LVs. There is no clear separation between BI-TBI and the combined set of BN-TBI and HC either along the x-axis (first LV) or the y-axis (second LV).

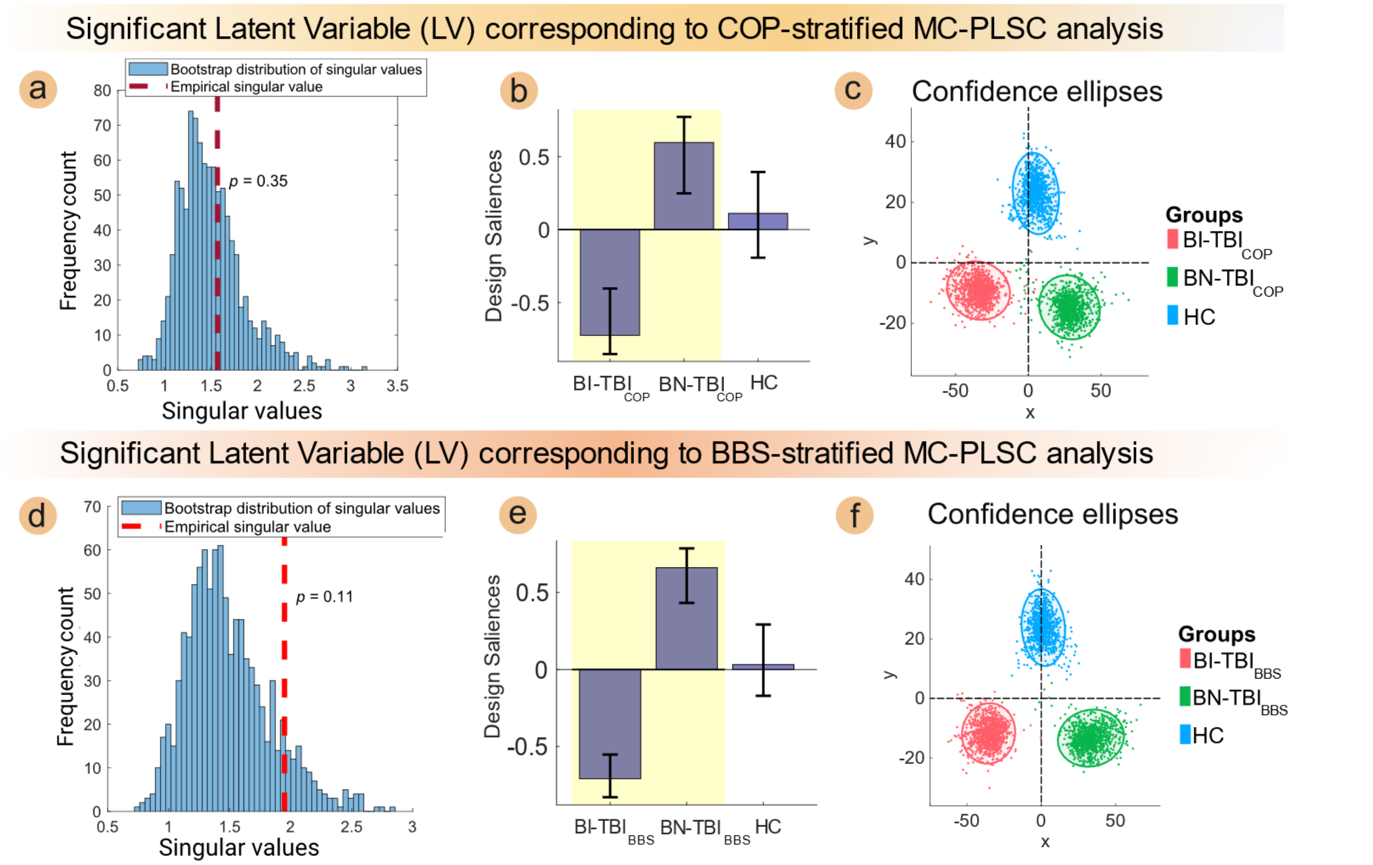


**Figure S5**: The results from the MC-PLSC, when implemented with the individuals stratified based on COP and BBS, are illustrated in the top and bottom row, respectively. The brain imaging variable is the **beta**-band weighted node degree, and the design variable is the group label. In (a) and (d), the empirical singular value associated with the first LV (red dotted line) is overlaid on the histogram of singular values (blue bar graph) computed from the permutation sample. In each instance, the empirical singular value is deemed statistically <u>not</u> significant since it remains below 95th percentile of the respective null distribution. The design salience **U** is represented by a bar plot in (b) and (e), where the height of each bar denotes the magnitude of an element of the first left singular vector. From (b) and (e), one can infer that BN-TBI is significantly different from BN-TBI, but neither of them from HC. The error bars generated with the respective singular vectors from the bootstrap sample are shown along with the bar plots to mark the 95% confidence interval. In both cases, BI-TBI and BN-TBI design saliences were robust (yellow background) as their bootstrap ratio was more than 2.5, but not that of HC. The 95% confidence ellipses of scatter plots in (c) and (f) are obtained with the first and second LVs. There is no clear separation between BI-TBI and the combined set of BN-TBI and HC either along the x-axis (first LV) or the y-axis (second LV).

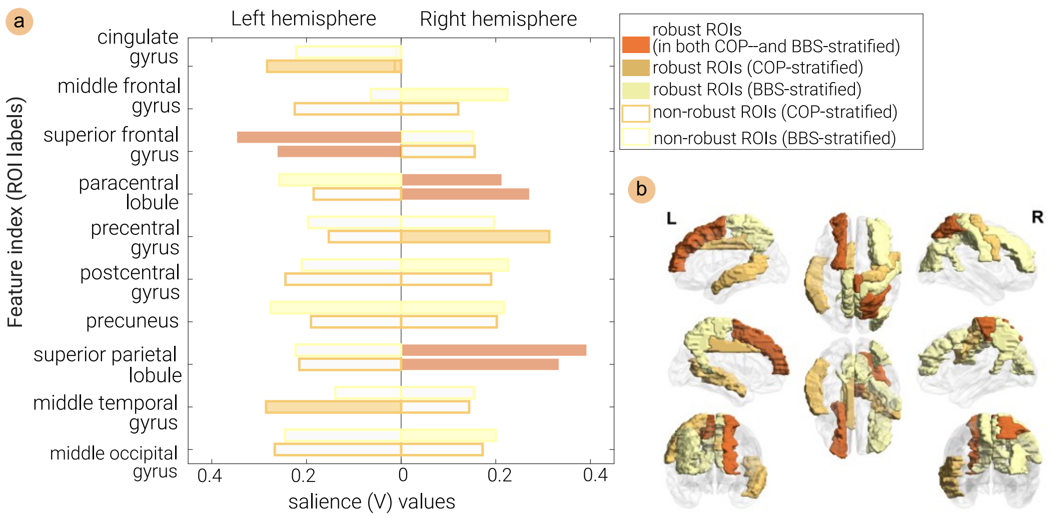


**Figure S6**: (a) The bar plot of brain saliences (functional connectivity strength of each ROI measured by the **alpha-band** imaginary part of coherence) associated with the significant LV. The top bar for each ROI denotes the magnitude of the brain salience for BBS-stratified analysis, whereas the bottom one corresponds to the COP-stratified analysis. The brain saliences were selected by the bootstrap test based on the criterion that BSR must exceed 2.5. (b) The diagrammatic representation of ROIs germane to stable alpha-band brain connectivity overlaid on the cortical surface. The functional connectivity strength of the left *superior frontal gyrus,* right *paracentral lobule* and *superior parietal lobule* were found to be robust to bootstrap testing in both COP- and BBS-stratified analysis, whereas the *right middle frontal gyrus, right middle occipital gyrus, bilateral precuneus, and left paracentral lobule* were robust to bootstrap testing in only BBS stratified analysis and *left middle temporal gyrus, right precentral gyrus,* and *cingulate gyrus* were robust only in COP-stratified analysis.

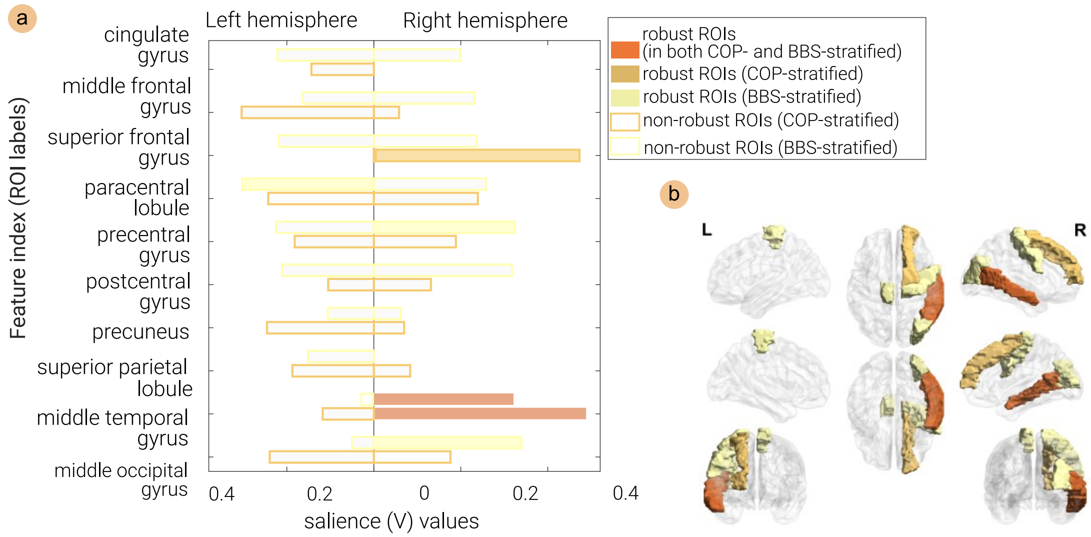


**Figure S7**: (a) The bar plot of brain saliences (functional connectivity strength of each region measured by the **beta-band** imaginary part of coherence) associated with the significant LV. The top bar for each ROI denotes the magnitude of the brain salience for BBS-stratified analysis, whereas the bottom one corresponds to the COP-stratified analysis. The brain saliences were selected by the bootstrap test based on the criterion that BSR must exceed 2.5. (b) The diagrammatic representation of ROIs germane to stable beta-band brain connectivity overlaid on the cortical surface. The functional connectivity strength of only the right *middle temporal gyrus* was found to be robust to bootstrap testing in both COP- and BBS-stratified analyses, whereas the *right superior frontal gyrus* was robust to bootstrap testing in only BBS stratified analysis.
